# Phylogenomic conflict coincides with rapid morphological innovation

**DOI:** 10.1101/2020.11.04.368902

**Authors:** Caroline Parins-Fukuchi, Gregory W. Stull, Stephen A. Smith

## Abstract

Evolutionary biologists have long been fascinated with the episodes of rapid phenotypic innovation that underlie the emergence of major lineages. Although our understanding of the environmental and ecological contexts of such episodes has steadily increased, it has remained unclear how population processes contribute to emergent macroevolutionary patterns. One insight gleaned from phylogenomics is that phylogenomic conflict, frequently caused by population-level processes, is often rampant during the origin of major lineages. With the understanding that phylogenomic conflict is often driven by complex population processes, we hypothesized that there may be a direct correspondence between areas of high conflict and elevated rates of phenotypic innovation if both patterns result from the same processes. We evaluated this hypothesis in six clades spanning vertebrates and plants. We found that the most conflict-rich regions of these six clades also tended to experience the highest rates of phenotypic innovation, suggesting that population processes shaping both phenotypic and genomic evolution may leave signatures at deep timescales. Closer examination of the biological significance of phylogenomic conflict may yield improved connections between micro- and macroevolution and increase our understanding of the processes that shape the origin of major lineages across the Tree of Life.

## Introduction

Phylogenomic conflict, where gene trees disagree about species tree resolution, is common across genomes and throughout the tree of life (Smith et al. 2015; Chen et al. 2017; Smith et al. 2020). Conflict is often driven by population processes such as incomplete lineage sorting (ILS), hybridization, and rapid evolution. Regions of high conflict often coincide with the emergence of major clades, such as mammals (e.g., Song et al. 2012), angiosperms (e.g., Wickett et al. 2014), and metazoa (e.g., Nosenko et al. 2013). This suggests that heterogeneity in the population processes underlying the origins of biodiversity has been recorded in particular nodes across the tree of life (e.g., Smith et al. 2020, OTPT Initiative 2019). Most large-scale phylogenetic and phylogenomic studies meant to resolve species relationships have treated gene-tree discordance as an analytical nuisance to be filtered or accommodated (e.g., Song et al. 2012; Wickett et al. 2014; OTPT Initiative 2019). However, since phylogenomic conflict often represents the imprint of past population genetic processes on the genome, studying its correlation with other macroevolutionary patterns may shed light on the microevolutionary processes underlying major transitions across the tree of life.

Researchers have long recognized that rates of morphological evolution vary across the tree of life, with pronounced bursts in morphological change interspersed with periods of relative stasis (Simpson 1944, Eldredge and Gould 1972). However, the strength of the relationship between rates of morphological change and genomic evolution is not well-understood. In particular, it may be worthwhile to examine whether phylogenomic conflict tends to coincide with periods of rapid morphological change. This may help to identify whether genomic conflict and morphological innovation detectable over macroevolutionary timescales stem from the same population processes.

Phylogenomic conflict often appears to coincide with important episodes of morphological differentiation among major lineages. For example, the major differences in life history and body plan that distinguish mammalian orders emerged rapidly among ancestral taxa following the K-Pg mass extinction (O’Leary et al. 2013). Phylogenetic resolution of many of the deepest mammal nodes has proved difficult and attempts at resolution have included the use of both sophisticated methodological approaches and the generation of large-scale genomic datasets (Morgan et al. 2013; Romiguier et al. 2013; Teeling et al. 2013). The early avian radiation has proved similarly challenging and is notable for the rapid establishment of phenotypically and ecologically disparate lineages. Several large-scale studies using massive genomic datasets have revealed extensive conflict among phylogenetic branches coinciding with the early radiation of crown Aves (Jarvis et al. 2014; Prum et al. 2015). Since the origin of land plants, there have been numerous major phases of morphological and ecological innovation, ranging from the initial appearance of vascular plant body plans in the late Silurian (Kenrick and Crane 1997) to distinct phases of angiosperm radiation from the Cretaceous to the present (Friis et al. 2011). As with the vertebrate lineages, the origins of many major plant clades show elevated levels of phylogenomic conflict, some of which may stem from biased representations of gene categories with functional significance (Lee et al. 2011). Many of these high-conflict branches have been the subject of intense analysis and debate (e.g., Drew et al. 2014; OTPT Initiative 2019; Smith et al. 2020) and recent sequencing efforts have failed to confidently resolve most of these challenges.

Here we examine whether periods of high genomic conflict coincide with episodes of rapid morphological evolution across several phylogenetically disparate clades. Many clades show qualitative evidence that time periods with particularly rampant conflict are associated with important phenotypic changes. However, if phylogenomic conflict indeed represents the remnants of complex population processes that are adaptively important, we reasoned that there should be a direct correspondence between areas of high conflict and elevated rates of phenotypic evolution. To examine whether this is the case, we reconstructed concurrent patterns in phylogenomic conflict and phenotypic evolution across six major clades of the eukaryotic tree of life: crown Aves, Mammalia, and the vascular plant lineages Ericales, Caryophyllales, Polypodiopsida, and Acrogymnospermae.

## Results

We assembled and analyzed six genomic datasets for plants, mammals, and birds. Specifically, we examined published phylogenomic datasets including 624 genes of Caryophyllales from Yang et al. (2018); 387 genes of Ericales from Larson et al. (2020); 1332 gene regions of Polypodiopsida from Shen et al. (2018); 1308 gene regions representing Acrogymnospermae from Ran et al. (2018); 424 genes of Mammalia from Song et al. (2012); and 259 gene regions of Aves from Prum et al. (2015). We gathered publicly available phenotypic matrices corresponding to each of these datasets (see Materials and Methods). For the Caryophyllales, a suitable matrix was not available, and so we generated a new matrix comprising 23 phenotypic traits scored across the species included in the phylogenomic sampling. Generally, broad phenotypic matrices that cover the taxa in each of these datasets are rare or non-existent. Furthermore, overlap between the genomic and phenotypic matrices was such that, other than for Caryophyllales, each needed to be reduced to ensure that identical edges were being compared. Divergence times continue to be controversial across the tree of life. To overcome these difficulties, we used one set of divergence times and one topology for both the morphological and molecular analyses (see Supplementary Information).

We found that mammals experienced elevated levels of both phylogenomic discordance and phenotypic innovation near the K-Pg boundary (Fig. 1A, 1B). These episodes of conflict coincided with high phenotypic rates calculated from two datasets: a massive ‘phenomic’ matrix of >4000 qualitative characters (O’Leary et al. 2013, median p=0.001; range=0-0.006) and a dataset of mammalian body masses (Jones et al. 2009, median p=0.144; range=0.117-0.175), which we interpreted as a proxy for many strongly correlated aspects of life history and ecology (Calder 1983). The two phenotypic datasets displayed remarkably similar patterns through time, showing that both life history and anatomical characteristics diversified rapidly during the early divergences of placental lineages. Although skeletal and dental characters comprised the majority of the qualitative dataset, characters representing internal and external organs, musculature, and behavior were also included. Our analysis of body masses are consistent with earlier studies in showing that higher-level variation in mammalian life history was established during their early radiation (Stearns 1983), coincident with rapid rates of anatomical change and high phylogenomic discordance. The association of body mass with conflict was statistically non-significant due to a continued elevated pace of body mass evolution after the K-Pg (Fig. 1B). While many of the phenotypic and ecological innovations of mammals likely originated during the evolutionary burst near the K-Pg boundary, it is possible that adjustments to body mass represent an additional source of more labile adaptive variation that has facilitated the continued expansion of niche-space occupied by mammalian lineages throughout the Cenozoic.

**Figure 1.**
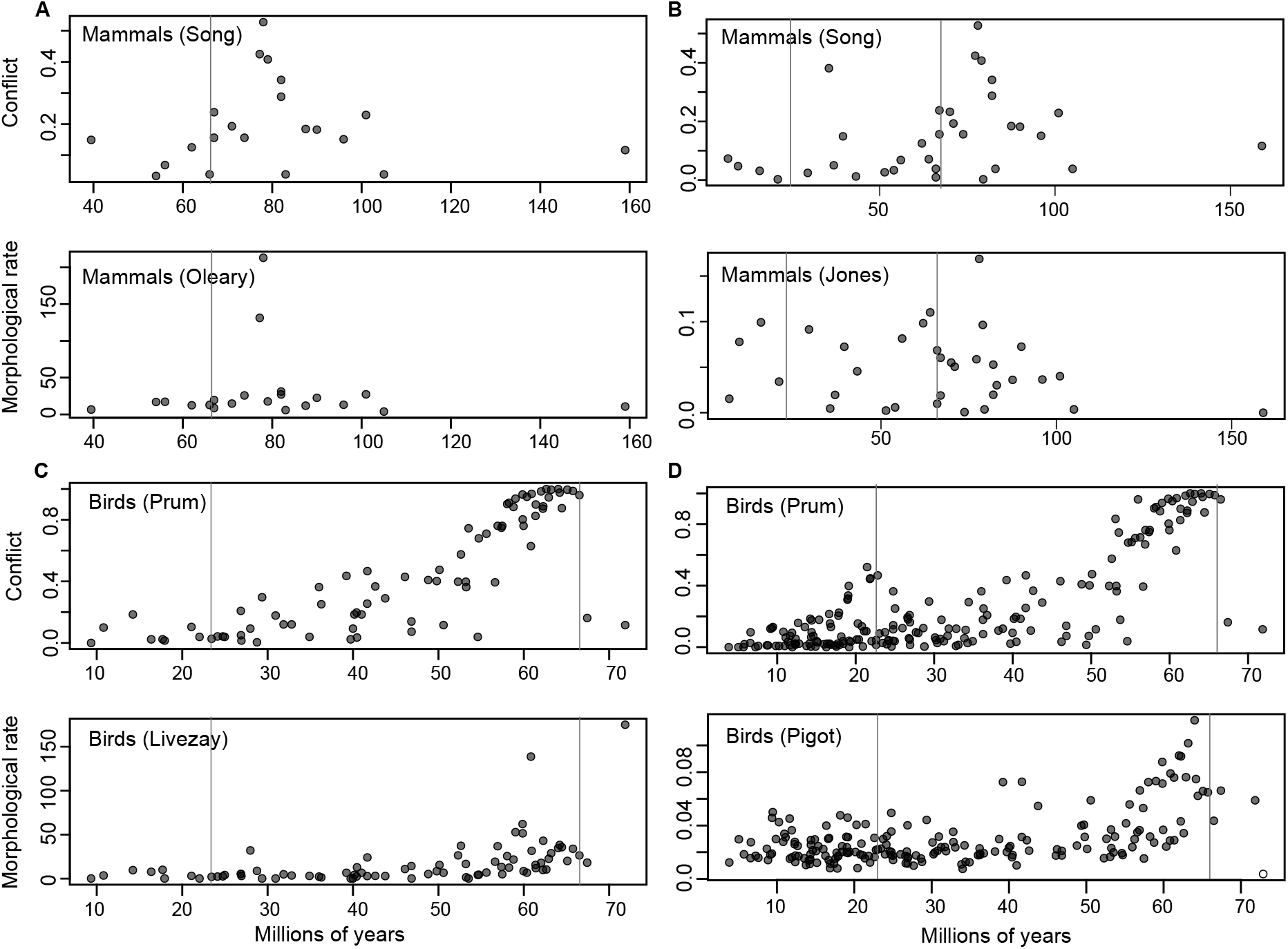
Co-occurring patterns in gene-tree conflict and morphological rates of evolution across the bird and mammal datasets. Conflict is measured as the proportion of gene trees that disagree with the species tree. Morphological rate is measured as the number of changes per million years for qualitative datasets and Brownain variance per million years for quantitative datasets. Vertical lines correspond to the onset of the Paleogene (66 Ma) and the Miocene (23.03 Ma).

For birds, our analyses of the genomic (Prum et al. 2015), qualitative (Livezay and Zusi 2007, median p=0; range=0-0.001), and quantitative (Pigot et al. 2019, median p=0; range=0-0.001) data revealed rampant conflict accompanied by elevated phenotypic rates among early-diverging avian lineages immediately following the K-Pg boundary (Fig. 1C, 1D). We also observed an additional episode of increased conflict that occurred at the onset of the Miocene. The results of the two phenotypic datasets were similar, with the main difference being that the quantitative dataset, comprised of principal component axes corresponding to body size, relative beak size, and relative wing size, showed slightly more elevated rates throughout the Cenozoic compared to the qualitative dataset, following the initial peak present in both datasets at the K-Pg. This difference likely reflects the increased lability of these three axes of quantitative variation versus the discrete transformations captured by the qualitative traits.

The Caryophyllales showed substantial conflict during epochs that also witnessed fast rates of phenotypic change (Fig. 2A; median p=0, range=0-0.004). Based on available divergence times, these conflicts occurred between ~59-101 MYA with the most extensive conflict at ~77 MYA. Caryophyllales clearly stand out among major flowering plant clades in terms of their phenotypic and ecological diversity (e.g., Brockington et al. 2009, 2011, 2013; Carlquist 2010; Ronse De Craene 2013; Ronse De Craene and Brockington 2013; Timoneda et al. 2019), and we find that many characteristic Caryophyllales features (e.g., betalain pigments, successive cambia, variations in perianth differentiation and merosity, campylotropous ovules, curved embryos, carnivory) arose during this temporal window of high conflict, with some traits showing complicated yet intriguing patterns of presence/absence across the phylogeny. Overall, the link between conflict and rate of innovation was very strong in the Caryophyllales, showing a close relationship both visually (Fig. 2A) and statistically (Fig. S2A).

**Figure 2.**
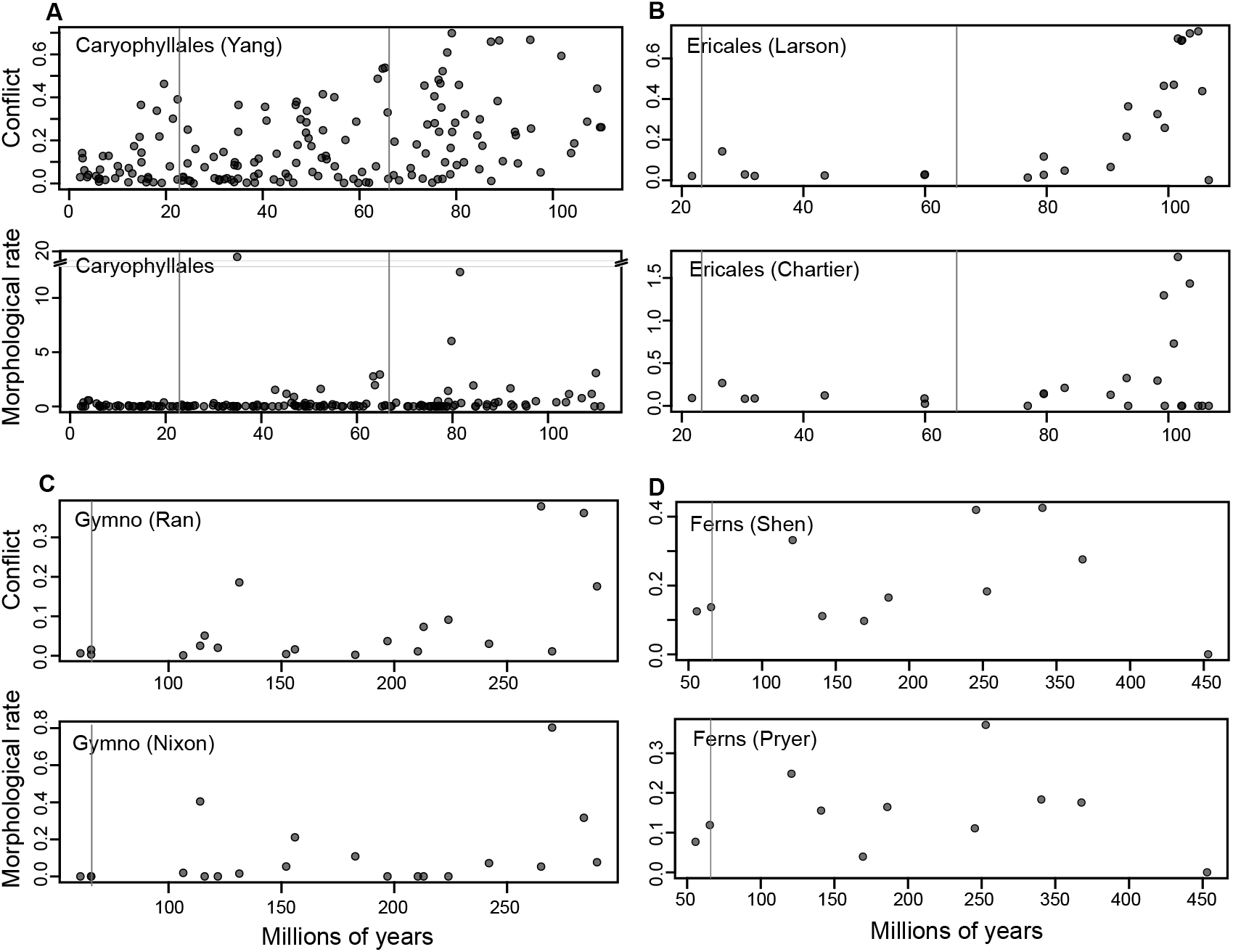
Co-occurring patterns in gene-tree conflict and morphological rates of evolution across major plant clades. Conflict is measured as the proportion of gene trees that disagree with the species tree. Morphological rate is measured as the number of changes per million years. Vertical lines correspond to the onset of the Paleogene (66 Ma) and the Miocene (23.03 Ma).

The Ericales exhibited both elevated conflict and rapid phenotypic change during their early radiation (Fig. 2B; median p=0.006, range=0.001-0.013). This episode witnessed the rapid evolution of reproductive features such as perianth merism, androecial structure (e.g., stamen number and anther orientation and attachment), and carpel number. Similar variation in floral morphology is evident in some of the earliest fossil representatives of this order (e.g., Schönenberger and Friis 2001; Crepet et al. 2013; Martínez et al. 2016) and likely relates to early co-evolution with insect pollinators, particularly hymenoptera (Crepet 1996; Nixon and Crepet 1993). This observed phylogenomic conflict and innovation in floral morphology precedes the evolution of the major families and also correspond to several life history shifts and movements out of the tropics (Rose et al. 2018) occurring by ~79-90 MYA based on available fossil evidence (e.g., Crepet et al. 2013) and between ~100-110 based on molecular divergence-time estimates (Magallón et al. 2015; Rose et al. 2018).

The gymnosperms (Acrogymnospermae) show several spikes in both conflict and rates of morphological evolution, loosely corresponding with major stratigraphic boundaries marked by episodes of major ecological upheaval (Fig. 2C; median p=0.8945, range=0.873-0.92). We found the visible similarity of the patterns in morphological rate and phylogenomic conflict compelling despite the lack of statistical significance and believe that the lack of significance likely stems from limited taxonomic sampling in both the morphological and molecular datasets. The largest spike for both conflict and innovation occurs before the P-T boundary and corresponds to the divergence of Gnetales. The phylogenetic placement of this lineage has long been contentious, with different molecular datasets (or analytical methods) resulting in different topologies (e.g., Zhong et al. 2010). Morphologically, Gnetales is also highly modified compared to other gymnosperms, which has led to various hypotheses concerning its placement (Crane 1985). Our results, which follow recent studies in placing Gnetales sister to Pinaceae (e.g., Ran et al. 2018; Smith et al. 2020), suggest that the divergence of the order involved a coordinated burst in both genomic and morphological evolution; the possible causes are unclear but deserve further study.

The analysis of ferns (=Polypodiopsida or ‘monilophytes’, Pryer et al. 2001) suggest a possible link between morphological rate and phylogenomic conflict that is qualitatively apparent but not statistically detectable (Fig. 2D; median p=0.534, range=0.504-0.573), perhaps due to the small sample size. Ferns (in the broad sense) are a heterogeneous assemblage of ancient lineages—including horsetails (Equisetiidae), marratioid ferns (Marattiidae), whisk and ophioglossid ferns (Ophioglossidae), and leptosporangiate ferns (Polypodiidae)—that trace back to the Devonian radiation of vascular plants, ca. 393 to 359 Ma (e.g., Kenrick and Crane 1997; Gensel et al. 2020). Prior to phylogenetic analyses including molecular data (e.g., Pyrer et al. 2001), the lineages comprising ferns s.l. were not thought to form a clade, but analyses have continued to recover this relationship with strong support (Pryer et al. 2001; Wickett et al. 2014; OTPT Initiative 2019). However, relationships among the four major constituent lineages remain contentious (e.g., Shen et al. 2018; OTPT Initiative 2019). Our results suggest that the conflict underlying earliest divergences in fern phylogeny may also coincide with pronounced phases of rapid morphological innovation, a possibility that deserves further study given that many aspects of phenotypic evolution in this clade—e.g., the origin(s) of megaphyll leaves—remain poorly known (Harrison and Morris 2017).

## Discussion

We demonstrate that several instances of high phylogenomic conflict correspond to instances of elevated rates in morphological innovation. This has implications for understanding both phenomena. First, biological processes, perhaps in addition to error, have a large impact on the patterns of conflict, even at deep time scales and when filtering for strongly supported signals. That biological processes can cause conflicting signals in gene trees is well known. However, typically, researchers attempt to overcome this conflict through modeling or methodological developments intended to construct a species tree. We suggest that population-level processes, such as population expansion or contraction during rapid speciation events, can leave strong macroevolutionary signatures observable in both patterns of gene-tree discordance and phenotypic evolution. The broad view engendered by our examination may be enhanced by targeted studies of the adaptive effects of ILS and other population-level processes during more recent radiations (e.g., Pease et al. 2016).

Disentangling the sources of conflict within phylogenomic datasets is fraught with difficulties. In the datasets examined here, while there may be instances of hybridization and, to a lesser extent, horizontal gene transfer that are contributing to the patterns of conflict, there is no published evidence that these phenomena are the source of biological conflict at the nodes that correspond to bursts in morphological innovation. There has been some suggestion of ancient hybridization early in the Caryophyllales (e.g., Morales-Briones et al. 2019); however, there are challenges of identifiability that have yet to be overcome. Whole genome duplications (perhaps in many cases stemming from hybridization, i.e., allopolyploidy events) may also result in conflicts across the tree of life, especially as they may create errors as a result of misidentified paralogy. Other systematic errors and background noise may be a source of conflict, especially when exacerbated by rapid evolution (Smith et al. 2020; Larson et al. 2020). Finally, incomplete lineage sorting has been recognized as a major source of gene-tree disagreement at all depths of the tree of life (Maddison and Knowles 2006). Conflicts due to incomplete lineage sorting will be amplified by particular population processes such as changes in effective population size from selection, bottlenecks, or other phenomena and/or rapid evolution (Pease and Hahn 2013). While errors also, undoubtedly, have contributed to some of the conflict we document, the vast majority of the nodes we highlight have been demonstrated to be difficult to resolve using multiple data types and analytical techniques (Jarvis et al. 2014; Prum et al. 2015; Wickett et al. 2015; Walker et al. 2017; OTPT Initiative 2019; Larson et al. 2020).

One notable attribute of several of the clades examined (e.g., Ericales, Caryophyllales) is that many phenotypic traits show complex patterns of presence/absence across lineages stemming from regions of high conflict. Such character-state distributions are typically interpreted as independent gains and losses (i.e., homoplasy). However, it is possible that the inheritance of phenotypic traits, from a polymorphic ancestral population, might in some cases follow an ILS-like process (i.e., hemiplasy; Avise and Robinson 2008; Guerrero and Hahn 2018). If the genes underlying phenotypic traits are subject to ILS, it follows that phenotypic traits themselves are also subject to this process. This might serve as a better explanation than independent gains/losses for complex distributions of phenotypic states in some clades. Hypothetically, this would imply that patterns of trait evolution during rapid “adaptive” radiations in some cases might be partly or largely stochastic—a byproduct of the sorting of underlying genetic variation. However, combinations of traits inherited through such a process could subsequently prove adaptive. This possibility deserves future attention and, importantly, would complement our results here. Elevated macroevolutionary rates of phenotypic evolution represent the rapid fixation of numerous traits within a short temporal window. This remains the case even if the substitution rate at the population level is artificially inflated due to unaccounted discordance (Mendes and Hahn 2016, Mendes et al. 2018). In cases of rapid speciation events from a polymorphic ancestor, hemiplasy might serve as one mechanistic link between genomic conflict and elevated rates of phenotypic evolution in the emergent lineages. Nevertheless, examining this possibility will be challenging given the complexity of the genotypic to phenotypic map and the noisiness of discrete morphological datasets.

A link between population processes and phenotypic innovation has long been theorized (Wright 1932). Although the specific relationship between microevolutionary processes and phenotypic changes observable at a macroevolutionary scale is not well-understood, previous work has considered how demographic events such as rapid fluctuations in population size and genic exchange between diverging populations might work synergistically with subsequent environmental adaptation to drive the exploration of diverse areas of morphospace by facilitating the emergence of both new phenotypic features and novel combinations of character states (Simpson 1944, Marques et al. 2019, Parins-Fukuchi 2020, McGee et al. 2020). Importantly, these processes are also likely to cause phylogenetic conflict if they persist over many generations and coincide with lineage divergences. We therefore suggest that phylogenomic conflict observable at deep timescales may often stem from the same population processes that facilitated the rapid and dramatic episodes of phenotypic change underpinning the origins of major clades. For example, the rapid speciation events that occur during evolutionary radiations may both generate conflicting phylogenomic signal due to population-level processes (e.g. rapid fluctuations in population size and selection) and drive the emergence of novel morphologies as diverging lineages carve out new ecological niches. In particular, a large population size will result in a smaller probability that two lineages will coalesce within a particular time. This should result in high conflict between gene trees. Several episodes of high conflict observed here (for example, in Ericales) are succeeded by periods of low conflict. This would suggest population size reduction if ILS can safely be assumed to be the source of conflict.

In addition to the notable coincidence of high conflict and rapid morphological innovation at particular nodes, many of these nodes also appear to correspond with major geologic events. As has long been observed, major radiations in mammals and crown birds seem closely tied to the K-Pg mass extinction; our results here linking conflict to morphological innovation hint at underlying processes that may have driven these radiations in the wake of K-Pg ecological collapse. Gymnosperms and ferns show coincidence spikes near the Permian-Triassic boundary, another widely studied mass extinction event (Benton and Twitchett 2003). But given our limited sampling of these particular clades, this association is much more tenuous. Unfortunately, examining these patterns in greater detail requires a reliance on accurate divergence times, which are challenging to estimate from molecular phylogenies. A better integration of fossil and phylogenomic data may offer a clearer understanding of how major geologic events have influenced coordinated bursts of conflict and phenotypic innovation.

### Conclusion

This study provides an important link between genomic and morphological evolution at deep timescales. We suggest that episodic evolutionary and population events leave signatures of conflict within genomes that may offer important insight on the processes responsible not only for conflict but also for massive changes in phenotype across disparate lineages. Moving forward, we suggest that closer examinations using more densely sampled molecular and morphological datasets representing taxa across the tree of life will help us to generate a deeper understanding of the micro- and macroevolutionary processes that drive major organismal changes from the genomic to the phenotypic level.

## Materials and Methods

### Genomic datasets

We examine several published phylogenomic datasets including 624 loci of Caryophyllales from Yang et al. (2018); 387 loci of Ericales from Larson et al. (2020); 1332 loci of ferns from Shen et al. 2018; 1308 loci of gymnosperms from Ran et al. (2018); 424 loci of mammals from Song et al. (2012); and 259 loci of Aves from Prum et al. (2015). Species trees were all based on those presented in the original publications. Gene trees were re-estimated using IQ-TREE (v. 1.6.11) with support calculated using the approximate likelihood ratio test (-alrt 1000) and with the GTR and Γ model for nucleotides (-m GTR+G) and the LG and Γ model for amino acids (-m LG+G).

### Morphological datasets

Large morphological datasets for major clades of flowering plants are surprisingly few and, additionally, most available datasets are relatively limited in their sampling of either taxa and/or characters. For the Carophyllales, we generated a new phenotypic dataset corresponding to the phylogenomic sampling, including 23 traits representing various vegetative and reproductive morphological features as well as several biochemical and physiological traits (e.g., betalain presence and photosynthetic pathway type). This matrix is available in the data supplement. For the gymnosperms, we used the published data from Nixon et al. (1994) where the data are primarily scored at family level. The taxon circumscriptions applied in Nixon et al. (1994) for their OTUs are monophyletic and consistent with our current understanding of gymnosperm phylogeny, and thus were applicable to the clades examined here. Thus multiple exemplars within a clade corresponding to one of their OTUs will be identical in their states. For the Ericales, we used the published matrix developed by Chartier et al. (2017) focused on floral characters. For mammals, we used the published dataset of O’Leary et al. (2013), which covers mammal morphology broadly, as well as body size data gathered from the panTHERIA database (Jones et al. 2009). For the ferns, we used the morphological dataset from Pryer et al. (2001), which was developed for analysis of fern relationships within the context of vascular plant phylogeny. Finally, for the birds, we used the morphological matrix compiled by Livezay and Zusi (2007) as well as the principal component scores calculated from the large dataset of linear measurements presented by Pigot et al. (2019). Many of the morphological datasets were sampled across more (or different) taxa than present in the genomic datasets—in these cases, we pruned the morphological datasets to include only the taxa present in the genomic data (and vice-versa).

### Phylogenomic analyses

Conflict analyses were conducted with the program *bp* (available from https://github.com/FePhyFoFum/gophy) using species and gene trees. We used the options -tv to print the concordance, conflict, and noninformative values back to the species tree and -scut 80 to ignore edges that had less than 80% for the aLRT statistic as calculated from the IQ-TREE analyses.

### Morphological analyses

We calculated rates of discrete character evolution by estimating the number of character transitions along each branch using parsimony and dividing this estimate by the amount of time represented by the corresponding branch in the time-scaled tree. Rates calculated from discrete-character matrices were therefore presented in units of inferred character transitions per unit time. For quantitative traits, we calculated the average character divergences along each branch by estimating ancestral states for each character under a Brownian motion model of evolution and then representing the amount of change along each edge as the absolute value of the difference between the trait value inferred for the ancestor and that inferred for the descendant. Like with the qualitative analyses, we then transformed these raw divergences to rates by calculating their ratio to the corresponding amount of elapsed time for the corresponding edge in the time-scaled tree. We also performed analyses on the discrete datasets using a model-based approach. We are concerned about the adequacy of existing Markov models for discrete phenotypic characters, but we provide these results in the supplement to provide further context for interested readers. Morphological branch lengths were calculated on the species tree topology using RAxML v8.2.12 (Stamatakis 2014) under the Mk model (-K MK) with ascertainment bias (-m ASC_MULTIGAMMA).

### Assessing statistical significance

We designed a permutation test to evaluate the statistical significance of the observed temporal associations between phylogenomic discordance and morphological rate. We removed all edges with either no conflict or no phenotypic change prior to performing the permutation test because such nodes increase in frequency as phylogenetic sampling increases. This skews the distribution heavily, causing nodes with only low or moderate conflict/rate to be erroneously identified as significantly high. We identified nodes with ‘significant’ elevated discordance and rate by resampling observed values 1000 times (with replacement) and generating an empirical null distribution using the means calculated for each replicate. Significant nodes were identified at a one-tailed 0.05 threshold. We then checked whether the number of nodes with both significantly high discordance and rate departed from the expectation if the same number of nodes was sampled randomly. This test examines the strength of the association between discordance and morphological rate, but was conservative for datasets with only a small number of nodes from which to sample (which yielded small overlap between high-discordance and high-rate nodes). Some degree of qualitative assessment using the discordance- and rate-through-time plots was therefore still needed.

### Selection of species tree topology

We used the same topology for both the morphological and molecular analyses. Conflict and morphological rates were mapped onto the species tree resolved in the study from which the genomic data were sourced.

## Data availability

Gene trees, dated species trees, phenotypic matrices, and code used to perform the analyses are available from Figshare (DOI: https://doi.org/10.6084/m9.figshare.13190816.v2).

## Acknowledgements and Funding

We thank J Pease for comments that improved the manuscript. CPF thanks G Auteri and M Foote for helpful conversations. SAS thanks his lab group for comments on early results. GWS thanks W DiMichele for comments on early results. CPF was supported as TC Chamberlin Postdoctoral Fellow in the Department of Geophysical Sciences at the University of Chicago during the conception and execution of this project. GWS acknowledges support from Chinese Academy of Sciences (CAS) President’s International Fellowship Initiative (No. 2020PB0009) and the China Postdoctoral Science Foundation (CPSF) International Postdoctoral Exchange Program. SAS was supported by National Science Foundation (NSF) grant 1917146.

